# Further evidence that CP-AMPARs are critically involved in synaptic tag and capture at hippocampal CA1 synapses

**DOI:** 10.1101/2020.07.28.224857

**Authors:** Pojeong Park, Heather Kang, John Georgiou, Min Zhuo, Bong-Kiun Kaang, Graham L. Collingridge

## Abstract

The synaptic tag and capture (STC) hypothesis provides an important theoretical basis for understanding the synaptic basis of associative learning. We recently provided pharmacological evidence that calcium-permeable AMPA receptors (CP-AMPARs) are a crucial component of this process. Here we have investigated two predictions that arise on the basis of CP-AMPARs serving as a trigger of the STC effect. Firstly, we compared the effects of the order in which we delivered a strong theta burst stimulation (TBS) protocol (75 pulses) and a weak TBS protocol (15 pulses) to two independent inputs. We only observed a significant STC effect when the strong preceded the weak TBS. Second, we found that pausing stimulation following either the sTBS or the wTBS for ∼20 min largely eliminates the STC effect. These observations are exactly as predicted for a process that is triggered by the synaptic insertion of CP-AMPARs and provide a framework for establishing the underlying molecular mechanism.

## INTRODUCTION

The concept of synaptic tag and capture (STC) was introduced to explain how input-specificity of synaptic plasticity could be maintained in the presence of *de novo* protein synthesis. In the initial pioneering experiments it was shown that a strong induction protocol to one input not only generated a protein-synthesis-dependent form of LTP at that input but was able to enable a subsequent weak induction protocol at an independent input to generate a larger and more sustained LTP (Frey and Morris, 1997). The original proposed mechanism is that a sufficiently strong induction protocol initiates cell-wide protein synthesis to generate plasticity-related proteins (PRPs) and that the weak protocol sets a synaptic tag that enables these locally-tagged synapses to capture PRPs. Since the discovery of this STC phenomenon there has been a massive effort to understand the underlying molecular mechanisms that are responsible for the formation of PRPs and the identity of the synaptic tag, given the relevance of this synaptic process to associative learning and memory (Redondo & Morris, 2011).

Previously, we showed that calcium-permeable (CP) AMPA receptors (CP-AMPARs) are transiently expressed during some forms of long-term potentiation (LTP) at Schaffer collateral-commissural synapses (Plant et al., 2006; Park et al., 2016) and proposed that these could be involved in the STC process. Recently, we provided direct evidence that this was indeed the case. We found that if we pharmacologically inhibited CP-AMPARs, during and shortly following the strong induction protocol, then the facilitation of LTP in response to the weak induction protocol was eliminated (Park et al., 2019). Therefore, we proposed that the synaptic activation of CP-AMPARs is involved in the triggering of PRPs, and that these newly synthesized proteins are engaged by the weak induction protocol to facilitate the LTP on the independent input. We also found that inhibiting CP-AMPARs during and following just the weak induction protocol resulted in a partial inhibition of the facilitation of LTP. We proposed therefore that CP-AMPARs are also required during induction of LTP by the weak input for the full heterosynaptic metaplastic effect to be observed.

This mechanism leads to two predications, that we have tested in the present study using theta burst stimulation (TBS) protocols. First, this CP-AMPAR-based mechanism requires the strong induction protocol to precede the weak one, since multiple trains are required to drive CP- AMPARs to the synapse under the recording conditions of our experiments. Second, on the assumption that once inserted CP-AMPARs need to be activated synaptically to elicit their effect, then stopping stimulation should mimic the effects of pharmacological inhibition of CP- AMPARs. To test these predications we interleaved four sets of experiments where the weak induction protocol comprised a single episode of TBS, termed weak TBS (wTBS; 15 stimuli), and the strong induction protocol comprised three episodes of TBS that were spaced in time with an inter-episode interval of 10 min, termed spaced TBS (sTBS; 75 stimuli). For set A, we delivered the wTBS 30 min after the sTBS, to replicate the control experiments reported previously (Park et al, 2019). For set B, we reversed the order of presentation of the sTBS and wTBS. For sets C and D, we reverted to sTBS before wTBS but stopped stimulation immediately following either the sTBS (set C) or the wTBS (set D). Our observations are entirely consistent with the notion that CP-AMPARs are inserted as a result of the sTBS to enable enhanced LTP to be induced on an independent input by the wTBS. They are also consistent with the idea that CP-AMPARs are additionally engaged by the wTBS to contribute to the induction of the facilitated LTP. Therefore CP-AMPARs are crucial for the STC process. They do so by initiating a heterosynaptic priming effect, where they locally trigger *de novo* protein synthesis, and they tag inputs for heterosynaptic metaplasticity.

## RESULTS

To compare the effects of the order of presentation of the weak and strong induction protocols we compared two protocols:

For set A, we delivered the sTBS 30 min before the wTBS was delivered to an independent input. As described previously for a different data set, this greatly facilitated the magnitude of LTP induced by the wTBS (Fig. 1), via a process that we refer to as heterosynaptic priming (Park et al., 2019). For example, the level of the LTP induced by a wTBS (averaged over a 1 min period, 90 min following TBS) was 124 ± 4%, n = 8 (Fig. 1A,D) and 156 ±7%, n = 11 (Fig. 1B,D) in the absence or presence of heterosynaptic priming (*p* = 0.0023, one-way ANOVA with Bonferroni’s *post hoc* test, F(2, 20) = 7.76). As noted previously (Park et al., 2019), the LTP induced by a sTBS, but not that induced by a wTBS, was also associated with a small heterosynaptic LTP (Fig. 1A, B).

**Fig 1.**
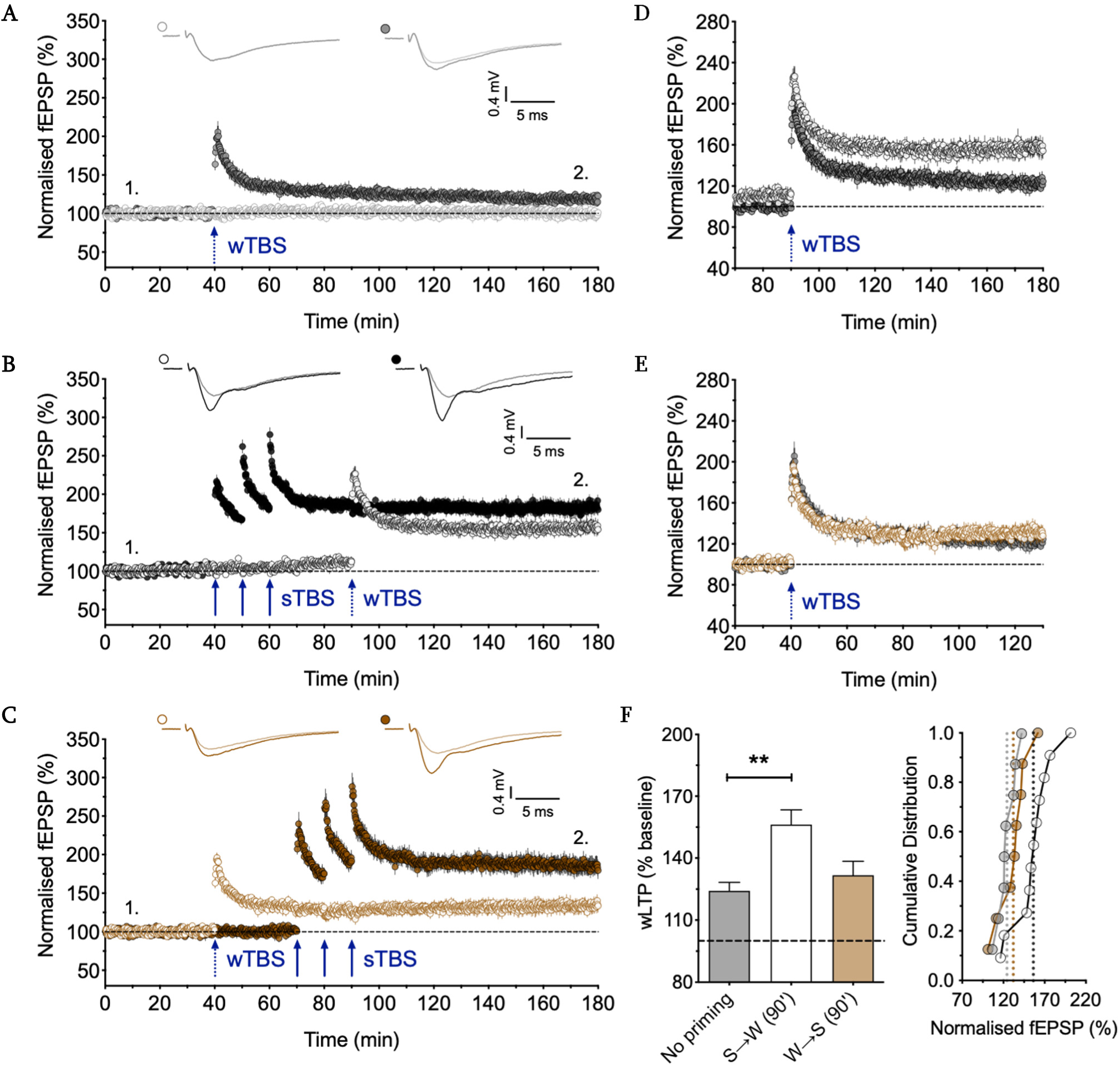
Evidence that sTBS before a wTBS more effectively engages the STC process. **A**. Two- input experiment that shows that a wTBS (15 pulses) induces a small, persistent input-specific LTP (n = 8). In this and subsequent time-course plots the data are mean ± SEM and representative traces show the superimposition of baseline and potentiated fEPSPs (averages of 5 successive records at the times indicated by numbers). **B**. Delivery of a sTBS to one input (filled circles) resulted in a modest heterosynaptic LTP in the second input plus an enhanced LTP induced by a wTBS (open symbols; n = 11). **C**. Delivery of a wTBS to one input (filled circles) resulted in a small LTP that was not appreciably affected by a subsequent sTBS on the other input (n = 8). **D**. Superimposition of the LTP induced by the wTBS illustrated in panels A and B. **E**. Superimposition of the LTP induced by the wTBS illustrated in panels A and C. **F**. Quantification of the LTP, 90 min post wTBS. \**p* < 0.05; \*\**p* < 0.01; \*\*\**p* < 0.001; comparisons vs. wTBS without heterosynaptic sTBS (no priming).

For set B, we reversed the order of presentation of the TBS (Fig. 1C, E). We found that the level of LTP induced by the wTBS was not significantly different whether a heterosynaptic priming stimulus was delivered (132 ± 7%, n = 8) or not (124 ± 4%, n = 8; *p* = 0.7370, one-way ANOVA with Bonferroni’s *post hoc* test). We noted a trend for an enhanced response (also see summary plots in Fig.1F), but this effect could be attributed to the small heterosynaptic LTP induced by the sTBS.

To test the effects of stopping stimulation we compared two additional protocols (Fig. 2). For set C, we stopped stimulation for 20 min following the third TBS episode (Fig. 2A, C); in addition, during the sTBS protocol we also stopped stimulation for most of the time following the second TBS episode. When compared to the experiments of Set A, we found that the level of LTP induced by the wTBS (118 ± 6%, n = 9) was effectively identical to that observed following the wTBS in the absence of heterosynaptic priming (Fig. 1D, 2F; *p* = 0.9716, one-way ANOVA with Bonferroni’s *post hoc* test) and was significantly less than that when heterosynaptic priming was employed (*p* = 0.0002, one-way ANOVA with Bonferroni’s *post hoc* test, F(2,25) = 12.13). Thus, interrupting stimulation during and following the sTBS completely prevented the STC process from occurring.

**Fig. 2.**
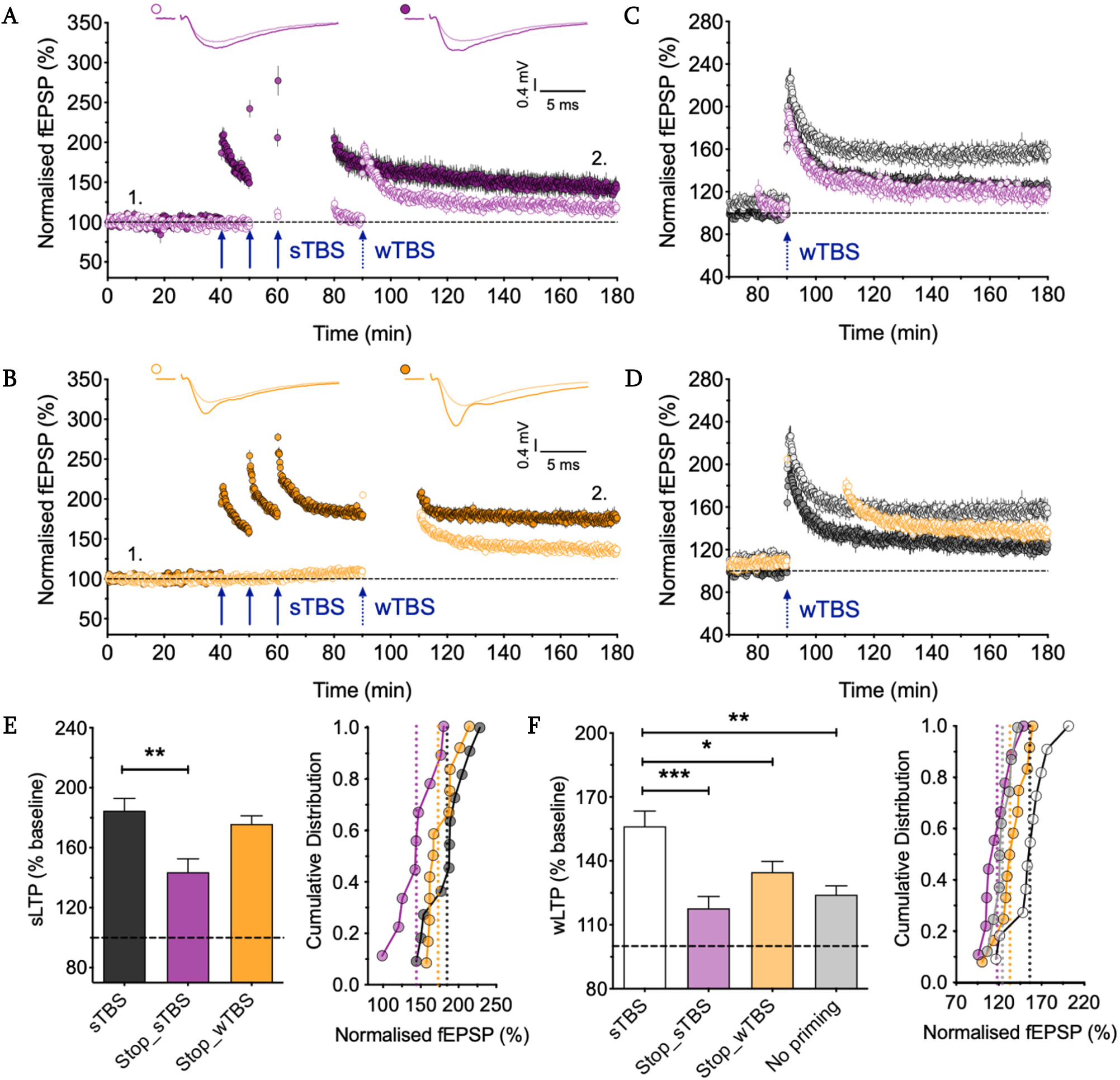
Evidence that stopping stimulation impairs the STC process. **A**. An experiment performed as in Fig. 1B, except that stimulation was paused on both inputs for 10 min following the 2nd TBS and 20 min following the 3rd TBS (n = 9). **B**. An experiment performed as in Fig. 1B, except that stimulation on both inputs was paused for 20 min following the wTBS (n = 12). **C**. Superimposition of the LTP induced by the wTBS illustrated in Fig. 2A, 1A, and 1B. **D**. Superimposition of the LTP induced by the wTBS illustrated in Fig.2B, Fig. 1A, and 1B. **E**. Quantification of the level of LTP induced by the sTBS for these experiments, 120 min post sTBS. **F**. Quantification of the level of LTP induced by the wTBS for these experiments, 90 min post wTBS. \**p* < 0.05; \*\**p* < 0.01; \*\*\**p* < 0.001; comparisons vs. sTBS.

**Fig. 3.**
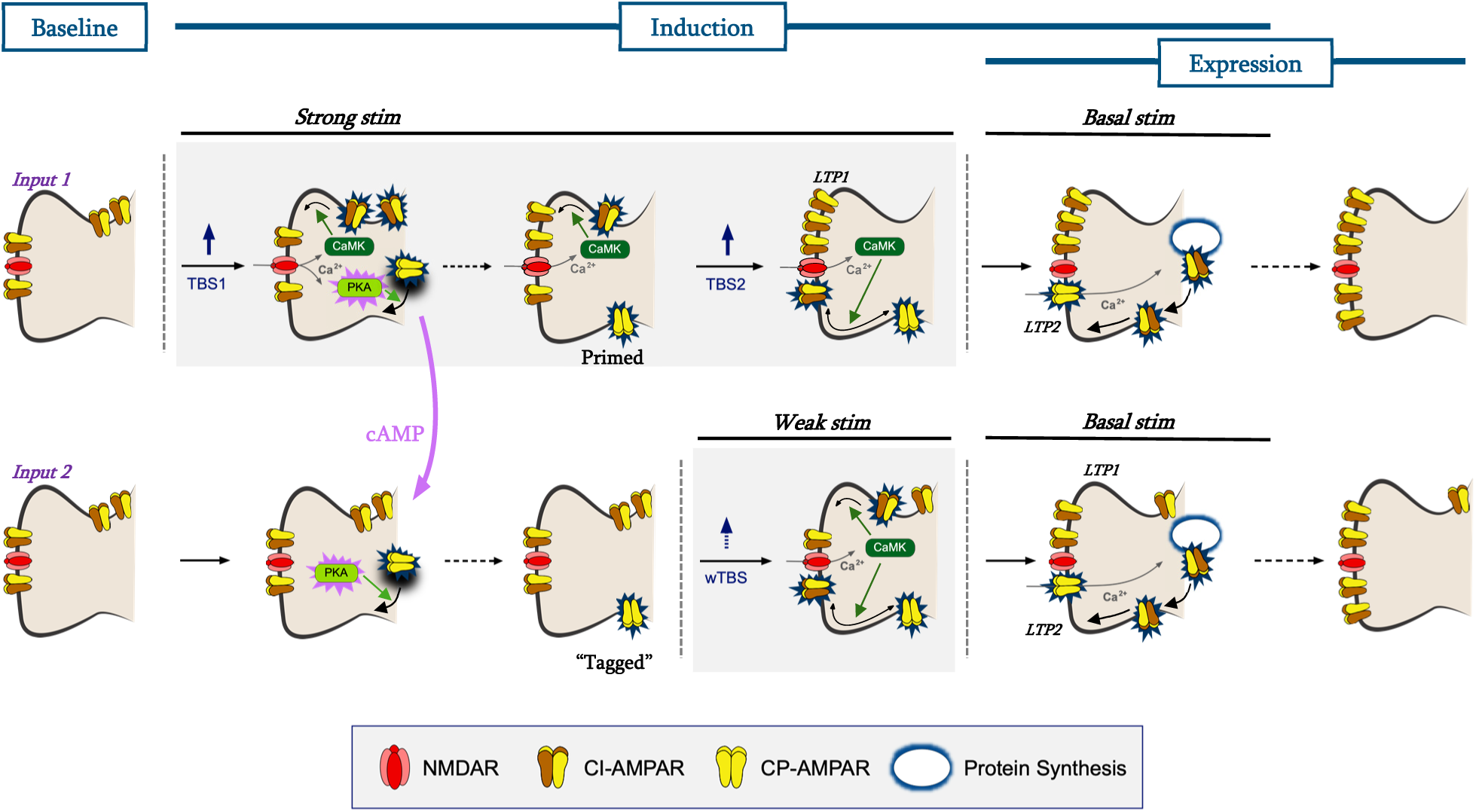
Hypothetical scheme for the STC process. Two inputs (shown in rows) are depicted by individual spine targets. The inputs are independent as defined by the lack of heterosynaptic paired-pulse facilitation. A strong stimulus comprised of spaced TBS (sTBS) is delivered to the upper input and a weak TBS (wTBS) stimulus (comprised of a short, single TBS) is delivered to the lower input, 30 min later. The first TBS, within the sTBS, activates NMDA receptors (NMDARs) to trigger homosynaptic LTP1 via engaging CaMKII to drive calcium-impermeable (CI-AMPARs) into the synapse. Simultaneously it primes for LTP2 by activating PKA, via the formation of cAMP, and this drives calcium-permeable (CP-AMPARs) onto the plasma membrane at perisynaptic sites. The subsequent TBS2 (and TBS3, not shown) drives these CP-AMPARs into the synapse where they increase the response amplitude by virtue of their higher single channel conductance. Basal stimulation activates these CP-AMPARs to drive *de novo* protein synthesis resulting in the insertion of additional CI-AMPARs at the expense of the transiently expressed CP-AMPARs. The cAMP diffuses to adjacent synapses where it activates PKA to drive CP-AMPARs into the plasma membrane. Some make it to the synapse where they trigger heterosynaptic LTP and may be activated in response to a wTBS to trigger LTP2 (not illustrated – see Park et al, 2019). Others are restricted to perisynaptic sites, where they “tag” these surround synapses for LTP2. A weak TBS delivered during the dwell time that these CP-AMPARs are present perisynaptically drives these CP-AMPARs into the synapse. Basal stimulation activates these CP-AMPARs to trigger local *de novo* protein synthesis and the generation of LTP2.

For set D, we stopped stimulation for 20 min following the wTBS episode. We found that the level of LTP induced by the wTBS following heterosynaptic priming (135 ±5 %, n = 12) was significantly less than that observed when there was no pause in stimulation following the wTBS (Fig 1B, D, Fig. 2F; *p* = 0.0215, one-way ANOVA with Bonferroni’s *post hoc* test) and not significantly different from that observed in the absence of heterosynaptic priming (Fig 1A, D, Fig. 2F; *p* = 0.4539, one-way ANOVA with Bonferroni’s *post hoc* test, F(2,28) = 7.39). The trend for enhanced LTP in this post-wTBS stop experiment could be explained by the small heterosynaptic potentiation. Therefore, interrupting stimulation following the wTBS also substantially reduced, if not abolished, the STC effect.

## DISCUSSION

In the present study we have extended our work that has investigated the hypothesis that CP-AMPARs are important for the STC (Plant et al., 2006; Park et al., 2019). We performed four interleaved sets of experiments (A-D) that support the notion that CP-AMPARs are critically involved in this process.

In set A, we confirmed our own observation, with an entirely new data set, that a wTBS delivered 30 min after a sTBS on an independent input leads to a greatly enhanced LTP compared to that induced by a wTBS alone. This result is fully in agreement with the original STC experiments of Frey & Morris (Frey and Morris, 1997). Previously we showed that it is the timing rather than the strength per se of the “strong” TBS that is critical for the STC effect (Park et al., 2019). We compared two protocols, each three episodes of theta (75 pulses in total) where the inter-episodes were either 10 s (compressed) or 10 min (spaced) and observed the STC effect only with the spaced protocol. Other work had shown that compressed and spaced LTP induction protocols differ in that the latter specifically engages a component of LTP that requires activation of PKA and *de novo* protein synthesis (e.g., Huang & Kandel, 1994) and CP-AMPARs (Park et al., 2018). In the present study we therefore used our standard sTBS protocol as the “strong” stimulus.

Since CP-AMPARs are only transiently inserted into these synapses, with a dwell time in the order of minutes (Park et al., 2020), we previously hypothesized (Plant et al., 2006) and then demonstrated (Park et al., 2019) that these receptors could serve to “tag” synapses for enhanced LTP. Our data supported a model whereby the sTBS initiated local *de novo* protein synthesis by leading to the transient insertion of CP-AMPARs in a two-step process that required, firstly, the insertion of these receptors at perisynaptic sites and, secondly, their movement into the synapse (Park et al., 2018). The first TBS in the episode solely engages the first step whereas the subsequent TBS episodes drive these newly plasma membrane inserted CP-AMPARs into the synapse. This mechanism can fully account for why spaced protocols are required to generate the protein-synthesis dependent component of LTP, that is commonly referred to as LTP2 (Bliss & Collingridge, 1993) or late-phase LTP (Huang & Kandel, 1994). Since the sTBS enables enhanced LTP at a heterosynaptic input, as defined by the lack of heterosynaptic paired-pulse facilitation, we referred to the process as heterosynaptic priming (Park et al, 2019), which is a form of heterosynaptic metaplasticity (Hulme et al, 2014).

### The order of presentation of “strong” and “weak” induction protocols

In set B, we examined the effects of delivery the wTBS before the sTBS. According to the original STC hypothesis this should also be effective, though less so than delivery of the sTBS beforehand (see Frey & Morris, 1997; 1998). However, according to our CP-AMPAR mechanism, the wTBS before the sTBS should not be effective. This is because a single episode of TBS, as in our wTBS protocol, would not be expected to drive CP-AMPARs into the synapse. Consistent with our hypothesis we did not observe a significant STC effect when we delivered the wTBS before the sTBS. We did, however, observe a trend for a facilitation of the subsequent LTP but we believe that this can be accounted for by the small heterosynaptic LTP that accompanies LTP induced by a sTBS (Frey & Morris, 1997; Park et al., 2019). Therefore, we can conclude that to establish the STC process the “strong” stimulus must precede the “weak” stimulus under our experimental conditions. Of course, this doesn’t discount the weak before strong protocol being effective under different circumstances. Although a single episode of TBS does not drive CP-AMPARs into synapses under the conditions of our experiments, there are conditions under which it can. For example, in the presence of rolipram, to inhibit breakdown of cAMP, a weak TBS effectively drives CP-AMPARs into synapses (Park et al., 2016). Also, neuromodulators, such as noradrenaline, enables a weak induction protocol to generate protein synthesis-dependent LTP (Connor et al., 2011; Zhang et al., 2013). Finally, acute stress can drive CP-AMPARs into synapses and thereby facilitate LTP (Whitehead et al., 2013). Indeed, the multiple ways of enhancing the synaptic insertion of CP-AMPARs likely has functional significance for neuromodulation. In summary, our new observations are fully consistent with the role of CP-AMPARs is triggering the STC process, since a sTBS is required to lead to the synaptic insertion of CP-AMPARs.

### The requirement for synaptic activation post TBS

In set C, we explored whether stopping stimulation for 20 min following the sTBS was sufficient to prevent the STC effect. The logic behind this experiment is: if basal (low frequency) stimulation is required to activate the newly inserted CP-AMPARs to drive Ca^2+^ into the synapse (Morita et al, 2014) then stopping stimulation should have the equivalent effect as applying a CP-AMPAR blocker, such as IEM-1460 (Park et al, 2019). This is indeed what we observed. For these experiments we also stopped stimulation following the delivery of the second TBS episode (apart from a few stimuli to monitor the level of potentiation) because these stimuli would be expected to activate the newly synaptically-inserted CP-AMPARs. The complete elimination of the STC effect by stopping stimulation re-enforces the essential role of CP- AMPARs in driving the process during the time window following the TBS protocol.

In set D, we asked whether stopping stimulation for 20 min following the wTBS was required for the STC effect. We again observed a significant effect, which is consistent with what we observed previously with IEM-1460. A partial inhibition of the STC effect was previously reported when an inhibitor of *de novo* protein synthesis is applied (Barco et al, 2002; Park et al., 2019). Therefore, we can conclude that for the full STC effect to be observed there has to be *de novo* protein synthesis triggered by CP-AMPARs that become synaptically available following the wTBS. The simplest explanation for this finding is that a sTBS not only drives CP-AMPARs homosynaptically but also drives CP-AMPARs heterosynaptically. In some cases, they may be inserted into the synapse where they mediate the small heterosynaptic LTP that is consistently observed when a protein synthesis-dependent LTP is induced. In other instances, they may dwell at perisynaptic sites awaiting a stimulus to drive them into the synapse, which the wTBS is able to do. In other words, they tag synapses for protein synthesis-dependent synaptic plasticity. Accordingly, a wTBS is able to drive these CP-AMPARs into the synapse where basal stimulation activates them to initiate highly localized protein synthesis. Appealing as this mechanism is, it is unlikely to be the entire explanation, since either stopping stimulation or the application of inhibitors of either CP-AMPARs or *de novo* protein synthesis only partially inhibits the heterosynaptic facilitation of LTP. At a subset of synapses, it is possible that the necessary activation of CP-AMPARs and *de novo* protein synthesis has already taken place and that the wTBS is just required to activate NMDA receptors for a different necessary component of the induction process. Further work will be required to establish more precisely the underlying mechanisms of these two components of the STC process.

### Multiple forms of LTP and the roles of multiple subtypes of glutamate receptors

The role of glutamate receptors in LTP at the Schaffer collateral-commissural pathway is complex. Initially, it was shown that NMDA receptors are required (Collingridge et al, 1983). Next it was found that mGluRs are sometimes necessary (Bortolotto et al, 1994). A role for CP-AMPARs was then identified in a KO mouse line, in which the GluA2 subunit was deleted (Jia et al, 1996). Finally, a role for kainate receptors was found at these synapses at an early developmental stage (Lauri et al, 2006). LTP is similarly complex, comprising a family of overlapping processes that all require the activation of NMDA receptors under most circumstances. There is an initial potentiation, termed short-term potentiation (STP), that decays as the pathway is stimulated (Volianskis et al., 2003). Then there are two forms of LTP that are stable over many hours (LTP1 and LTP2) that are distinguished on the basis of their requirement for *de novo* protein synthesis (Bliss et al., 2018). Longer lasting forms of LTP also require transcription (LTP3). Here we have focused on LTP1 and LTP2. We have provided additional evidence that CP-AMPARs confer heterosynaptic metaplasticity by priming synapses for enhanced LTP (i.e., converting LTP1 into LTP2 at the primed synapses). This process is therefore distinct from the role of mGluRs in LTP at these synapses where they are involved in a metaplasticity that is entirely homosynaptic in nature (Bortolotto et al, 1994), though again involves *de novo* protein synthesis (Raymond et al, 2000).

As originally hypothesised, the STC process is an ideal candidate synaptic mechanism for long-lasting associative memory (Frey & Morris, 1997). Initially it was assumed that the *de novo* protein synthesis occurred at the soma and hence a tag was required to capture PRPs at synapses to enable their amplification. Subsequently, it was assumed that protein synthesis may occur locally (Redondo & Morris, 2011). Our mechanism (Park et al., 2019) is based on local protein synthesis. Accordingly, the role of the tag is not to capture PRPs *per se*, but to mark a surround of synapses that are able to undergo enhanced LTP by enabling an LTP1-inducing stimulus to generate LTP2. This mechanism is entirely consistent with the notion of clustered synaptic plasticity (Govindarajan et al, 2011).

### Concluding Remarks

In summary, we can conclude that CP-AMPARs are an integral part of the STC process. They are inserted into the synapse in two stages, firstly via an NMDA receptor-triggered PKA-dependent insertion into extrasynaptic sites and then by an NMDA receptor-triggered CaMKII-dependent movement into the synapse. The former provides many opportunities for modulation via neurotransmitters and other factors that regulate cAMP levels in neurons. It also enables a heterosynaptic nature to synaptic plasticity, since CP-AMPARs may be inserted at sites outside of the activated synapses. The dwell time of CP-AMPARs on the plasma membrane is in the order of minutes so that they can associate signals that are appropriately spaced in time.

## METHODS

Experiments were performed as described in Park et al. (2019). Briefly, transverse hippocampal slices (400 μm) were prepared from male C57BL/6 mice (10-12 weeks of age) using a vibratome (Leica, VT1200S). The CA3 region was cut, with a scalpel blade, to suppress the upstream neuronal excitability, and the slices were transferred to an incubation chamber that contained the recording solution (artificial cerebrospinal fluid, ACSF; mM): 124 NaCl, 3 KCl, 26 NaHCO3, 1.25 NaH2PO4, 2 MgSO4, 2 CaCl2 and 10 D-glucose (carbonated with 95% O2 and 5% CO2). Slices were allowed to recover at 32-34°C for 30 min, and then maintained at 26-28 °C for a minimum of 1 h before recordings were made. Hippocampal slices were continuously perfused at 3-4 mL/min with the oxygenated ACSF at 32°C. Two bipolar stimulating electrodes were positioned in stratum radiatum on either side of the recording electrode at approximately the same distance from the cell body layer. Two independent Schaffer collateral-commissural pathways (SCCPs) were stimulated alternately to obtain the evoked synaptic responses, each at a frequency of 0.1 Hz. The initial slope of evoked fEPSPs (V/s) was monitored and analysed using WinLTP (Anderson and Collingridge, 2007). Following a stable baseline period of at least 40 min, LTP was induced using theta-burst stimulation (TBS) delivered at the same basal stimulus intensity and pulse width (0.1 ms; constant voltage stimulator). An episode of TBS comprised bursts of 5 pulses at 100 Hz delivered at 5 Hz. The wTBS consisted of one episode of 3 bursts (i.e., 15 pulses in total). The sTBS comprised 3 episodes, each of 5 bursts, delivered with an inter-episode interval of 10 min (i.e., 75 pulses in total). Each experiment was performed on a slice obtained from a different mouse; therefore, n values denote both numbers of slices and mice used. Representative sample traces are an average of 5 consecutive responses, collected from typical experiments (stimulus artefacts were blanked for clarity). All four groups (A-D) were interleaved. Data are presented as mean ± SEM (standard error of the mean). Responses were normalised to the baseline prior to LTP induction, and data are expressed as % baseline. Statistical significance was assessed using one-way ANOVA with Bonferroni’s correction; the level of significance is denoted on the figures as follows: \**p* < 0.05, \*\**p* < 0.01 and \*\*\**p* < 0.001.

## ACKNOWLEDGEMENTS

This work was supported by the CIHR (GLC), the EJLB-CIHR Michael Smith Chair in Neurosciences and Mental Health, Canada Research Chair, and Canadian Institute for Health Research operating grants (CIHR66975 and 84256) (MZ) and the National Honor Scientist Program of the National Research Foundation funded by the South Korea Government (B-KK). This work was also supported by the Brain Canada Foundation through the Canada Brain Research Fund, with the financial support of Health Canada.

